# The lateral periaqeductal gray and its role in controlling the opposite behavioral choices of predatory hunting and social defense

**DOI:** 10.1101/2020.09.02.273961

**Authors:** Ignacio Javier Marín-Blasco, Miguel José Rangel, Marcus Vinicius C. Baldo, Simone Cristina Motta, Newton Sabino Canteras

**Affiliations:** Dept. Anatomy, Institute of Biomedical Sciences, University of São Paulo; São Paulo, SP 05508-000, Brazil; Dept. Physiology and Biophysics, Institute of Biomedical Sciences, University of São Paulo; São Paulo, SP 05508-000, Brazil

**Keywords:** prey hunting, social agonistic behavior, motivation

## Abstract

Evasion from imminent threats and prey attack are opposite behavioral choices critical to survival. Curiously, the lateral periaqueductal gray (LPAG) has been implicated in driving both responses. The LPAG responds to social threats and prey hunting while also drives predatory attacks and active defense. However, the LPAG neural mechanisms mediating these behaviors remain poorly defined. Here, we investigate how the LPAG mediates the choices of predatory hunting and evasion from a social threat. Pharmacogenetic inhibition in Fos DD-Cre mice of neurons responsive specifically to insect predation (IP) or social defeat (SD) revealed that distinct neuronal populations in the LPAG drive the prey hunting and evasion from social threats. We show that the LPAG provides massive glutamatergic projection to the lateral hypothalamic area (LHA). Optogenetic inhibition of the LPAG-LHA pathway impaired IP but did not alter escape/attack ratio during SD. We also found that pharmacogenetic inhibition of LHA^GABA^ neurons impaired IP, but did not change evasion during SD. The results suggest that the LPAG control over evasion to a social attack may be regarded as a stereotyped response depending probably on glutamatergic descending projections. On the other hand, the LPAG control over predatory behavior involves an ascending glutamatergic pathway to the LHA that likely influences LHA^GABA^ neurons driving predatory attack and prey consumption. The LPAG-LHA path supposedly provides an emotional drive for prey hunting and, of relevance, may conceivably have more widespread control on the motivational drive to seek other appetitive rewards.

## Introduction

The periaqueductal gray (PAG) has been commonly recognized as a downstream site in neural networks for the expression of a variety of behaviors, i.e., sexual, maternal, and defensive behaviors with accompanying modulation of nociceptive transmission, autonomic changes, and vocalization (Bandler and Shipley, 1994; Besson et al., 1991; Fanselow, 1991; Gruber-Dujardin, 2010; Jurgens, 1994; Lonstein and Stern, 1997, 1998; Lovick, 1993; Sakuma and Paff, 1979). By and large, PAG-related responses have been regarded as being mostly stereotyped and dependent on descending projections to the brainstem and spinal cord. However, the PAG is also known to influence complex events like approach and avoidance responses to perform risk assessment of potential threats (Cezario et al, 2008; Sukikara et al., 2010; Souza and Carobrez, 2016; Pobbe et al., 2011), fear memory processing (Deng et al., 2016; Di Scala et al., 1987; Di Scala and Sandner, 1989; Kincheski et al., 2012; Kim et al., 2013; de Andrade Rufino et al., 2019), and reward-seeking (Comoli et al., 2003; Sukikara et al., 2006; Mota-Ortiz et al., 2012; Tryon and Mizumori, 2018).

It is particularly puzzling that the PAG controls both aversive and rewarding behaviors. In this regard, we call special attention to the LPAG that has been implicated in mediating the opposite behavioral choices of predatory attack and evasion from a conspecific aggressor. Previous studies from our laboratory have associated the LPAG to the organization of prey hunting. Insect predation was associated with increased Fos expression in the LPAG (Comoli et al., 2003). Behavioral observations have shown that LPAG NMDA lesions interfere with prey hunting; the animals lost their motivation to pursue and attack prey without affecting the general levels of arousal, locomotor activity, and regular feeding (Mota-Ortiz et al., 2012). In line with these observations, the LPAG contains reward excited neurons, which exhibited an increased firing rate in response to appetitive food (Tryon and Mizumori, 2018). Conversely, the LPAG has also been shown to respond to a conspecific aggressor. Thus, a significant Fos upregulation in the LPAG was found in animals exposed to dominant conspecifics (Motta et al., 2009). In this regard, chemical and optogenetic stimulation experiments suggest that the LPAG mediates active defensive responses such as circa-strike defense, and supports a role in active social defensive behaviors to escape from a dominant’s attacks (Depaulis et al. 1992; Assareh et al. 2016; Wang et al. 2019).

Here, we investigated how the LPAG controls the opposite behavioral choices of predatory hunting and evasion from a social attack. We started asking whether there is a differential activation of the neural populations in the LPAG activated during insect predation (IP) and social defeat (SD). To this end, we used an approach based on the temporal translational dynamics of c-*fos* gene combining immunofluorescence for Fos protein and fluorescent *in situ* hybridization for c-*fos* mRNA (Fos protein/c-*fos* mRNA IF-FISH) in animals exposed sequentially to IP and SD. We next examined the functional role of the neural populations in the LPAG responsive to IP or SD and used pharmacogenetic silencing in Fos DD-Cre mice (Dillingham et al. 2019), which allowed targeting neural populations activated by a specific stimulus. Further, considering that LHA is one of the main targets of the LPAG (Mota-Ortiz et al. 2012), we used optogenetic inhibition and examined how LPAG projections to the lateral hypothalamic area (LHA) influence predatory and social defensive behaviors.

The LHA, in its turn, is involved in controlling predatory hunting and evasion from imminent threats (Li et al. 2018). Considering that the LHA is heavily targeted by the LPAG, we also investigated how the LHA mediates IP and SD. First, we differentiated neuronal populations in LHA responding to IP and SD using Fos protein/c-*fos* mRNA IF-FISH. Next, we determined the GABAergic and glutamatergic nature of LHA neurons responding to IP and SD using Fos protein/*VGAT* mRNA and Fos protein/*VGLUT2* mRNA IF-FISH. Finally, using VGAT-IRES-Cre and VGlut2-IRES-Cre mice, we applied pharmacogenetic tools to silence the activity of the LHA^GABA^ and LHA^Glut^ neurons selectively and then explored the functional role of LHA^GABA^ and LHA^Glut^ neurons on predatory hunting and social defensive behavior.

We here show that the LPAG contains two segregated neural populations that separately control opposite behavioral choices of predatory hunting and evasion from a social attack. Our findings are in line with the idea that the PAG works as a unique hub driving stereotyped responses and supplying primal emotional tone to influence complex aversive and appetitive responses (see Motta et al., 2017). Our results show that LPAG control over evasion in response to a social attack may be regarded as a stereotyped response depending probably on descending projections to brainstem sites organizing the behavioral output. On the other hand, the LPAG control over predatory behavior involves an ascending glutamatergic pathway to the LHA that likely influences LHA^GABA^ neurons to drive predatory hunting. This path supposedly exerts a role in providing an emotional drive for prey hunting and may conceivably have more widespread control on the motivational drive to seek other appetitive rewards.

## Results

### Identification of neuronal populations in LPAG activated by IP and SD

To differentiate neuronal populations in LPAG responding to insect predation (IP) and social defeat (SD), we combined immunofluorescence for Fos protein (IF) and fluorescent *in situ* hybridization for c-*fos* mRNA (FISH) following the protocol described by Marin-Blasco et al. (2017). Exposure to a first treatment increases c-*fos* mRNA levels, which reach a maximum level at about 20 min and decline to the basal levels two hours later when the Fos protein is close to its maximum (Kovács and Sawchenko, 1996; Zangenehpour and Chaudhuri, 2002; Kovács, 2008). The animals are then exposed to a second stimulus, and a second peak on c-*fos* mRNA will be observed nearly 20 min later. Thus, neurons responding to the first stimulus will show Fos protein (IF+/FISH−) mainly, neurons responding to the second (SD) will show mostly c-*fos* mRNA (IF−/FISH+), and those neurons activated by both stimuli will appear as double-labeled (IF+/FISH+). To determine the optimum exposure and sacrificing time points, we quantified the number of Fos protein- and c-*fos* mRNA positive-neurons in LPAG at different time points after IP and SD (see Figure S1). As shown in Figure S1, two hours after the set of IP, we found maximum levels of Fos protein levels and imperceptible c-*fos* mRNA signal; we obtained maximum c-*fos* mRNA levels 20 minutes after exposure to SD. However, low but perceptible newly synthesized Fos protein emerged at this time point in response to SD. Thus, we decided to shorten this time point to 15 minutes when c-*fos* mRNA levels are close to the maximum; less significant Fos protein levels were then detected (Figure S1). Since exposure to a stressing stimulus such as SD affects animals’ motivation for hunting, we exposed animals first to IP and subsequently to SD. We also included two control groups of animals exposed twice to the same stimulus (groups IP+IP and SD+SD) to discard neuronal activation due to the mere manipulation of animals. Animals repeatedly exposed to the same stimulus showed reactivation of about 50% of neurons (IF+/FISH+ neurons) and a low number of IF-/FISH+ neurons (Figure 1A,C). Animals exposed sequentially to IP and SD showed reactivation of about 40% of neurons (IF+/FISH+) in addition to a significantly higher number of newly activated neurons (IF−/FISH+) compared to IP+IP and SD+SD groups (Figure 1C).

**Figure 1.**
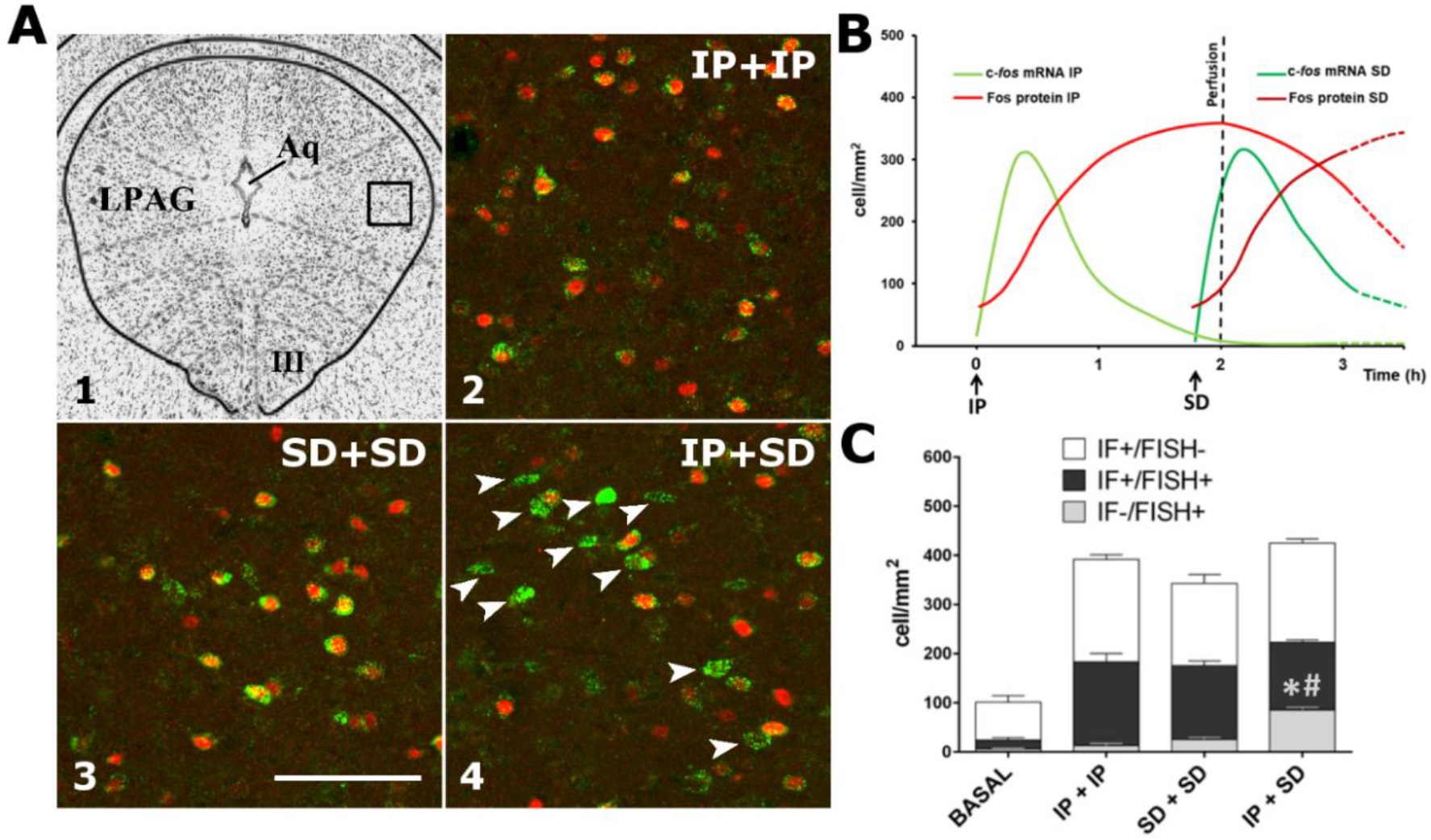
Identification of neuronal populations in LPAG activated by IP and SD. **(A) A1**. Photomicrograph of a representative transverse thionin-stained section of the LPAG at the level of the oculomotor nucleus (III). Square-delineated region indicates the approximate location of higher magnification fluorescence photomicrographs showing Fos/c-*fos* mRNA double labeling in (A2), (A3) and (A4). Aq – cerebral aqueduct. **A2-A4**. Representative Fos/c-*fos* mRNA double labeling fluorescence photomicrographs of groups IP+IP **(A2)**, SD+SD **(A3)**, and IP+SD **(A4)**. Fos protein positive cells are labeled by immunofluorescence in red (Alexa 594) and c-*fos* mRNA positive cells are labeled by fluorescent *in situ* hybridization in green (FITC). Arrows indicate newly activated neurons (IF−/FISH+) in response to the second stimulus (SD). Scale bar, 100 μm. **(B)** Temporal dynamics of c-*fos* mRNA and Fos protein in response to IP and SD (see **Supplementary Fig. 1**). Note that the prefusion occurred 15 min after the set of SD, when c-*fos* mRNA levels are close to the maximum and no significant Fos protein levels in response to SD were detected. **(C)** Median values of the density of IF+/FISH-, IF−/FISH+, and IF+/FISH+ neurons across the groups. GzLM analysis of the number of newly activated neurons (IF−/FISH+) in animals exposed sequentially to IP and SD showed a significant effect of group factor (groups IP+IP, SD+SD, and IP+SD) [χ^2^(2) = 165.57, *p* < 0.001]. Group IP+SD presented a higher number of newly activated neurons (IF−/FISH+) when compared to IP+IP (**\*)** and SD+SD (**#)** groups (*p* < 0.001). Groups: BASAL (*n* = 4); IP+IP (*n* = 4); SD+SD (*n* = 4); IP+SD (*n* = 6). Data are reported as mean ± SEM.

It is important to note that we would have the maximum levels of c-*fos* mRNA 20 minutes after the second stimulus (SD). However, we decided to sacrifice the animals 5 minutes earlier to minimize the detection of newly synthesized Fos protein. Thus, in this account, the number of newly activated neurons (IF−/FISH+) could have been underestimated. Despite this issue, the results suggest that exposure to SD induced activation of neurons already recruited by IP and further led to the recruitment of new neurons. Therefore, LPAG neuronal populations responding to predatory hunting and social defensive responses appear to be partially differentiated.

### Functional analysis of neuronal populations in LPAG activated by IP and SD

In Fos DD-Cre mice, Cre recombinase is fused to an *E. coli* dihydrofolate reductase (ecDHFR)-derived destabilizing domain (DD-Cre; Iwamoto et al. 2010; Sando et al., 2013) and is expressed under the control of the c-*fos* promoter. The DD-Cre construct is unstable and degraded via proteasome in the absence of the ecDHFR inhibitor trimethoprim (TMP). The intraperitoneal administration of TMP makes Cre recombinase stable (protected from degradation) and catalytically active (Figure 2A).

**Figure 2.**
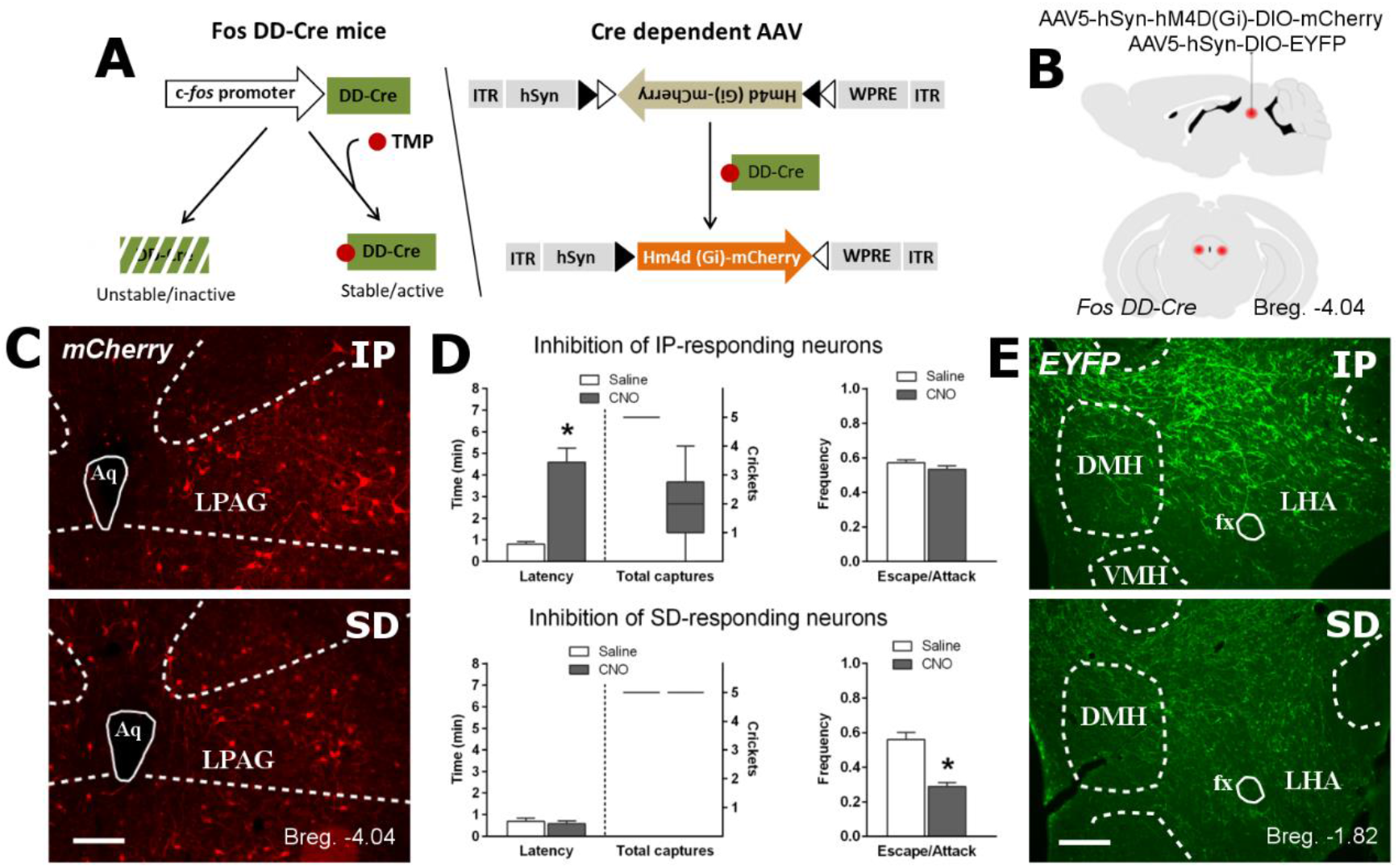
Pharmacogenetic inhibition of IP and SD-responding LPAG-neuronal populations. **(A)** Strategy for manipulation of active neurons in Fos DD-Cre mice. Administration of TMP induces stabilization of Cre recombinase expressed in active neurons thus promoting recombination in AAV vectors and expression of genes of interest – hM4D(Gi) inhibitory DREADD fused to mCherry and and EYFP. **(B)** Schematics showing the location of the bilateral AAV viral vectors injection in the LPAG of Fos DD-Cre mice. **(C)** Fluorescence photomicrographs from representative cases showing mCherry expressing neurons activated in response to IP (top) and SD (bottom). Aq – cerebral aqueduct. Scale bar, 150 μm. **(D)** The effect of CNO-induced inhibition of IP- (top) and SD-responding (bottom) neurons on predatory (latency to catch the first prey and total number of captures; left) and social defensive (escape/attack ratio; right) behaviors. The data for latency and escape/attack ratio are shown as mean ± SEM, and the data for the number crickets captured are represented as box plot graphs. Groups: IP Cre stabilized animals (*n* = 7); SD Cre stabilized animals (*n* = 7). For the latency to catch the first prey, Univariate ANOVA revealed a significant interaction between the factors (time dependent Cre stabilization x Treatment; F1,21=43,78; p<0.001). Post hoc pairwise comparisons (Tukey’s HSD test) revealed that IP Cre stabilized animals treated with CNO presented a significant increase in the latency to start hunting (p<0.001). IP Cre stabilized treated with CNO also presented a reduced number of captured crickets. For the escape/attack ratio, univariate ANOVA revealed a significant interaction between the factors (time dependent Cre stabilization x Treatment; F1,21=30.8; p<0.001). Post hoc pairwise comparisons (Tukey’s HSD test) revealed that SD Cre stabilized animals treated with CNO presented a significant decrease in the escape/attack ratio observed during the social agonistic interaction (p<0.001). **(E)** Fluorescence photomicrographs from representative cases showing the distribution of projection to the LHA from LPAG EYFP expressing neurons activated during IP (top) and SD (bottom). Projections originating from IP-responding neurons were substantially denser than those originating from SP-responding neurons. DMH – dorsomedial hypothalamic nucleus; fx – fornix; LHA – lateral hypothalamic area; VMH – ventromedial hypothalamic nucleus. Scale bar, 150 μm. **See S3 - videos**.

Here, Fos DD-Cre mice received bilateral injections in the LPAG of a Cre-dependent adeno-associated viral vector (AAV) expressing hM4D(Gi) inhibitory DREADD (Designer Receptor Exclusively Activated by Designer Drug) fused to mCherry reporter (AAV5-hSyn-hM4D(Gi)-DIO-mCherry) to selectively silence the activity of the LPAG neurons responsive to IP or SD (Figure 2A,B). Previous tract-tracing studies showed that LPAG neurons send substantial ascending projections to the LHA (Mota-Ortiz et al. 2012). Therefore, we also aimed to detect possible differences in the LHA projection patterns from LPAG-activated neurons in response to IP or SD. To this end, animals also received a paired injection of Cre-dependent AAV expressing EYFP reporter (AAV5-hSyn-DIO-EYFP) for tracing ascending projections to the lateral hypothalamic area (LHA) (Figure 2B). The EYFP reporter was used to trace ascending projections because it yields a much stronger fluorescent signal than the mCherry fused to hM4D(Gi). The administration of TMP in animals previously exposed to IP or SD stabilizes DD-Cre expressed in active neurons and leads to Cre-dependent expression of hM4D(Gi) and EYFP (Figure 2A). Animals were subsequently treated with saline or clozepine-N-oxide (CNO, Armbruster et al., 2007) and re-exposed to IP and SD for behavioral testing.

Previous studies showed that cytotoxic lesions in the LPAG increased the latency to start hunting and decreased the number of captured prey (Mota-Ortiz et al. 2012). Therefore, during predatory hunting, we quantified the latency to catch the first cricket and the total number of crickets captured. Conversely, chemical and optogenetic stimulation experiments suggest that the LPAG mediates predominantly active defensive responses such as the circa-strike defense. This supports a role in active social defensive behaviors aiming to escape from resident’s attacks (Depaulis et al. 1992; Assareh et al. 2016; Wang et al. 2019). Thus, during the social agonistic interaction, we analyzed the number of intruder escapes in response to the resident attacks (escape/attack ratio) as a measure of active defensive behavior.

Our results revealed that CNO treatment of animals with Cre stabilized for IP presented a significant increase in the latency to start hunting (p<0.001) (Figure 2D). In contrast, CNO-treated animals that received TMP Cre stabilization for SD did not differ from the saline-treated animals in the latency to start hunting (p>0.26) (Figure 2D). Likewise, in comparison to the other groups, CNO-treated animals that received TMP Cre stabilization for IP also presented a reduced number of captured preys (Figure 2D). Conversely, Cre-stabilized animals for SD that received CNO presented a significant decrease in the escape/attack ratio observed during the social agonistic interaction (p<0.001) (Figure 2D). In contrast, CNO-treated animals that received TMP Cre stabilization for IP did not differ from the saline-treated animals in the escape/attack ratio observed during SD (p>0.46) (Figure 2D).

Thus, our functional analysis in Fos DD-Cre mice revealed that CNO inhibition of LPAG neurons activated in response to IP led to an increase in the latency to start hunting and a decrease in the total number of crickets captured but did not alter escape/attack ratio during SD. Conversely, CNO inhibition of LPAG SD-responding neurons did not change predatory behavior parameters and led to a decrease in the escape/attack ratio during SD.

The results of our tracing analysis in Fos DD-Cre mice revealed that LPAG IP-responding neurons yielded a much denser EYFP anterograde labeling in LHA that those found for the animals that received TMP time-dependent Cre stabilization during SD (Figure 2E).

### Functional analysis of LHA-projecting LPAG terminals during IP and SD

The LPAG projects densely to the LHA, which is one of the main targets of the LPAG (Mota-Ortiz et al. 2012). We next investigated whether the LPAG – LHA projection influences predatory and social defensive behaviors. Considering the predominantly excitatory nature of this projection (see Supplementary Material 2 and Figure S2), we employed a projection-based optogenetic silencing approach to examine the effect of the inhibition of the LPAG – LHA projection on the predatory hunting and social defense. For LPAG – LHA projection photoinhibition, we bilaterally injected the LPAG of C57BL/6 mice with adeno-associated viral (AAV) vectors encoding Archaerhodopsin 3 (eArch3)—a trafficking-enhanced light-sensitive (589 nm) proton pump (Mattis et al. 2012) fused with enhanced yellow fluorescence protein (AAV5-hSyn-eArch3.0-EYFP) (Figure 3B). We included a control group of animals injected with a virus not expressing eArch3.0 (AAV-hSyn-EYFP-PA) to discard light stimulation behavioral effects. The 561-nm laser light was continually delivered to the LHA during the 5 min of IP or SD through surgically implanted dual-fiber optic elements.

**Figure 3.**
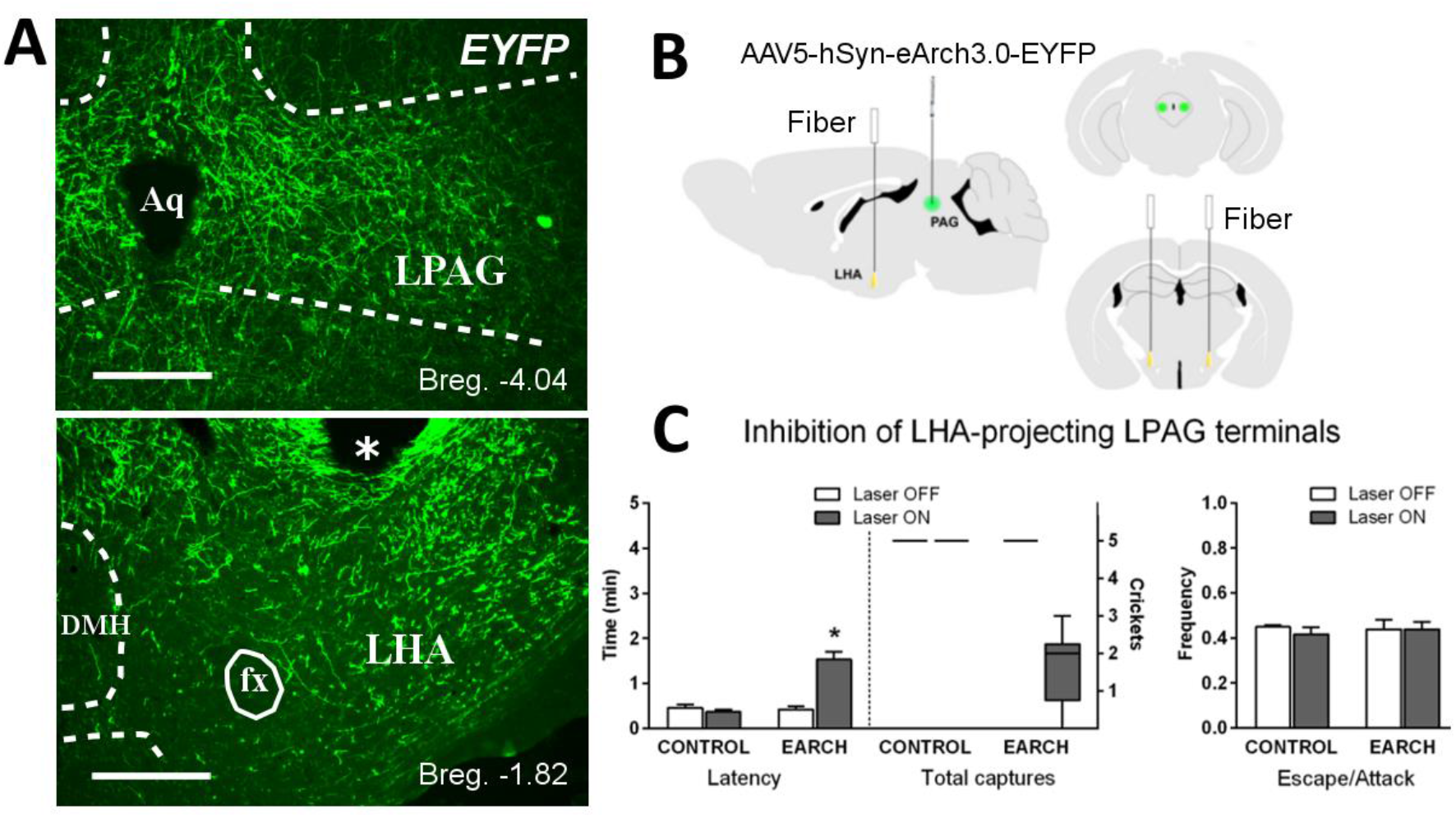
Optogenetic inhibition of LPAG-projecting LHA terminals in *C57BL/6 mice*. **(A)** Fluorescence photomicrographs illustrating the viral injection site in the LPAG with neurons expressing EYFP (top) and EYFP labeled fibers in LHA originating from LPAG (bottom). **(*)** tip of the optic fiber. Aq – cerebral aqueduct; DMH – dorsomedial hypothalamic nucleus; fx – fornix; LHA – lateral hypothalamic area. Scale bars, 200 μm (LPAG) and 150 μm (LHA). **(B)** Schematics showing the location of the bilateral AAV viral vectors injection in the LPAG and the position of bilateral optical fibers implanted in the LHA. **(C)** The effect of optogenetic inhibition of LPAG-projecting LHA terminals on predatory (latency to catch the first prey and total number of captures; left) and social defensive (escape/attack ratio; right) behaviors. The data for latency and escape/attack ratio are shown as mean ± SEM, and the data for the number crickets captured are represented as box plot graphs. Groups: CONTROL (*n* = 4) and EARCH (*n* = 6). For the latency to catch the first prey, univariate ANOVA revealed a significant effect for the factor virus (eArch+ and control, F1,8=21.28; p<0.002) and the factor treatment (laser on / off, F1,8=16.87; p<0.004), and a significant interaction between the factors (Virus x Treatment, F1,8=32.21; p<0.001). Post hoc pairwise comparisons (Tukey’s HSD test) revealed that eArch+ animals during laser on presented a significant increase in the latency to catch the first prey (p<0.001). Photoinhibition of the LPAG – LHA projection during IP also reduced the number of captured crickets. For the escape/attack ratio, univariate ANOVA revealed no significant effect for the factors virus (F1,8=0.53; p=0.48) and treatment (F1,8=0.21; p=0.659). **See S4-Videos**.

Our functional analysis revealed that photoinhibition of LPAG -LHA pathway impaired IP and increased the latency to start hunting while decreasing the total number of captured crickets; it did not alter escape/attack ratio during SD (Figure 3C).

### Identification of neuronal populations in LHA activated by IP and SD

To differentiate neuronal populations in LHA responding to IP and SD, we next applied IF-FISH for Fos protein and c-*fos* mRNA (Marin-Blasco et al., 2017). Animals repeatedly exposed to the same stimulus (groups IP+IP and SD+SD) showed reactivation of about 40% of neurons (IF+/FISH+ neurons) and an exceedingly low number of IF−/FISH+ neurons (Figure 4F,G). Animals exposed sequentially to IP and SD showed reactivation of about 38% of neurons (IF+/FISH+) in addition to a significantly higher number of newly activated neurons (IF−/FISH+) compared to IP+IP and SD+SD groups (Figure 4F,G). Therefore, the LHA neuronal populations responding to predatory hunting and social defensive responses appear to be partially differentiated in the LPAG.

**Figure 4.**
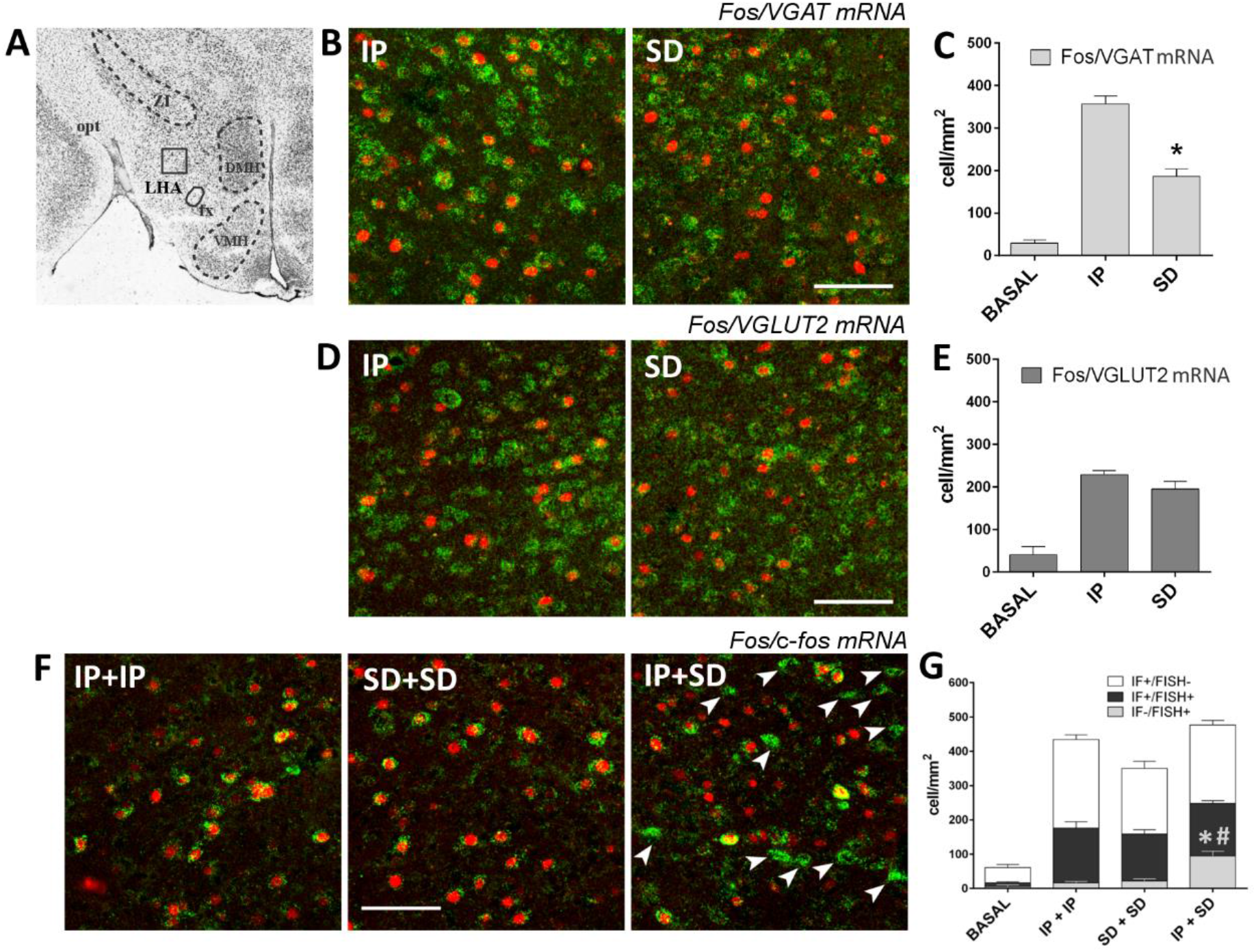
Analysis of the GABAergic and glutamatergic nature of LHA neurons recruited in response to predatory hunting and social defensive behavior and identification of LHA neural populations activated by IP and SD. **(A)** Photomicrograph of a representative transverse thionin-stained section at tuberal levels of the LHA. Square-delineated region indicates the approximate location of higher magnification fluorescence photomicrographs showing Fos/VGAT and VGLUT2 mRNAs double labeling in (B) and (D). DMH – dorsomedial hypothalamic nucleus; fx – fornix; LHA – lateral hypothalamic area; opt – optic tract; ZI zona incerta. **(B)** Representative fluorescence photomicrographs illustrating *Fos/VGAT* mRNA double labeling in the LHA of animals exposed to IP and SD. Fos protein positive cells are labeled by immunofluorescence in red (Alexa 594) and *VGAT* mRNA positive cells are labeled by fluorescent *in situ* hybridization in green (FITC). Scale bar, 75 μm. **(C)** Median values of the density of Fos/*VGAT*mRNA double labeled cells in the BASAL (*n* = 3), IP (*n* = 6), and SD (*n* = 6) groups. GzLM analysis of the number of Fos/*VGAT* mRNA neurons showed a significant effect of GROUP factor (groups Basal, IP, and SD) [χ^2^(2) = 197.71, *p* < 0.001]. SD group presented a lower number of Fos/*VGAT* mRNA neurons when compared to IP group (\**p* < 0.001). **(D)** Representative fluorescence photomicrographs illustrating *Fos/VGULT2* mRNA double labeling in LPAG of animals exposed to IP and SD. Fos protein positive cells are labeled by immunofluorescence in red (Alexa 594) and *VGLUT2* mRNA positive cells are labeled by fluorescent *in situ* hybridization in green (FITC). Scale bar, 75 μm. **(E)** Median values of the density of Fos/*VGLUT2*mRNA double labeled cells in the BASAL (*n* = 3), IP (*n* = 6), and SD (*n* = 6) groups. GzLM analysis of the number of Fos/*VGLUT2* mRNA neurons showed a significant effect of group factor [χ^2^(2) = 86.75, *p* < 0.001]. IP and SD groups presented an equivalent number of Fos/*VGLUT2* mRNA neurons. **(F)** Representative Fos/c-*fos* mRNA double labeling fluorescence photomicrographs of groups IP+IP, SD+SD, and IP+SD. Fos protein positive cells are labeled by immunofluorescence in red (Alexa 594) and c-*fos* mRNA positive cells are labeled by fluorescent *in situ* hybridization in green (FITC). Arrows indicate newly activated neurons (IF−/FISH+) in response to the second stimulus (SD). Scale bar, 75 μm. **(G)** Median values of the density of IF+/FISH−, IF−/FISH+, and IF+/FISH+ neurons in the BASAL (*n* = 4), IP+IP (*n* = 4), SD+SD (*n* = 4), and IP+SD (*n* = 6) groups. GzLM analysis of the number of newly activated neurons (IF–/FISH+) showed a significant effect of group factor (groups IP+IP, SD+SD, and IP+SD) [χ^2^(2) = 36.17, *p* < 0.001]. Control groups IP+IP and SD+SD did not differ in the number of new neurons. Group IP+SD presented a higher number of newly activated neurons (IF−/FISH+) when compared to IP+IP (**\***) and SD+SD (**#**) groups (*p* < 0.001). Data are reported as mean ± SEM.

### Characterization of GABAergic and glutamatergic LHA neurons responding to IP and SD

Recent optogenetic experiments suggest an essential role of LHA GABAergic neurons (LHA^GABA^) in driving predatory hunting (Li et al. 2018) as well as other motivated behaviors related to feeding (Jennings et al., 2013, 2015; O’Connor et al., 2015). Conversely, optogenetic stimulation of LHA glutamatergic neurons (LHA^Glut^) induces a robust aversive response in the real-time place-preference assay (Lazaridis et al. 2019); LHA^Glut^ neurons increase their activity during evasion but not during food chasing (Li et al. 2018). To characterize the GABAergic and glutamatergic nature of the LHA neurons activated by IP and SD, we next combined immunofluorescence for Fos protein and fluorescent *in situ* hybridization for mRNAs of the vesicular transporters of GABA (*VGAT*) and glutamate (*VGLUT2*), respectively (Figure 4B,D). The results showed that the number of Fos/*VGAT* mRNA double-labeled neurons was significantly higher after exposure to IP versus SD (Figure 4C) whereas the number of Fos/*VGLT2* mRNA double-labeled neurons was comparable in both conditions (Figure 4E). This suggested more significant recruitment of LHA^GABA^ neurons during predatory hunting perhaps reflecting a more important role of LHA^GABA^ neurons in IP. Conversely, activation of LHA^Glut^ neurons was similar for both IP and SD. This did not infer a differential role of the LHA^Glut^ neurons in any one of these behaviors.

### Functional role of LHA^GABA^ and LHA^Glut^ neurons on predatory hunting and social defensive behavior

To address the behavioral role of LHA^GABA^ and LHA^Glut^ neurons on predatory hunting and social defensive responses, we used a Cre-dependent AAV vector expressing an inhibitory DREADD (AAV5-hSyn-H=hM4D(Gi)-DIO-mCherry in VGAT-IRES-Cre and VGlut2-IRES-Cre mice to silence the activity of the LHA^GABA^ and LHA^Glut^ neurons selectively (Figure 5A,B). We also included a group of control animals injected with a Cre-dependent AAV not expressing the inhibitory DREADD (AAV5-hSyn-DIO-EYFP). Our functional analysis revealed that pharmacogenetic inhibition of LHA^GABA^ neurons impaired IP increasing the latency to start hunting and decreasing the total number of captured crickets, but it did not alter the escape/attack ratio during SD (Figure 5C). Conversely, pharmacogenetic inhibition of LHA^Glut^ neurons decreased the escape/attack ratio during SD but did not influence IP (Figure 5D).

**Figure 5.**
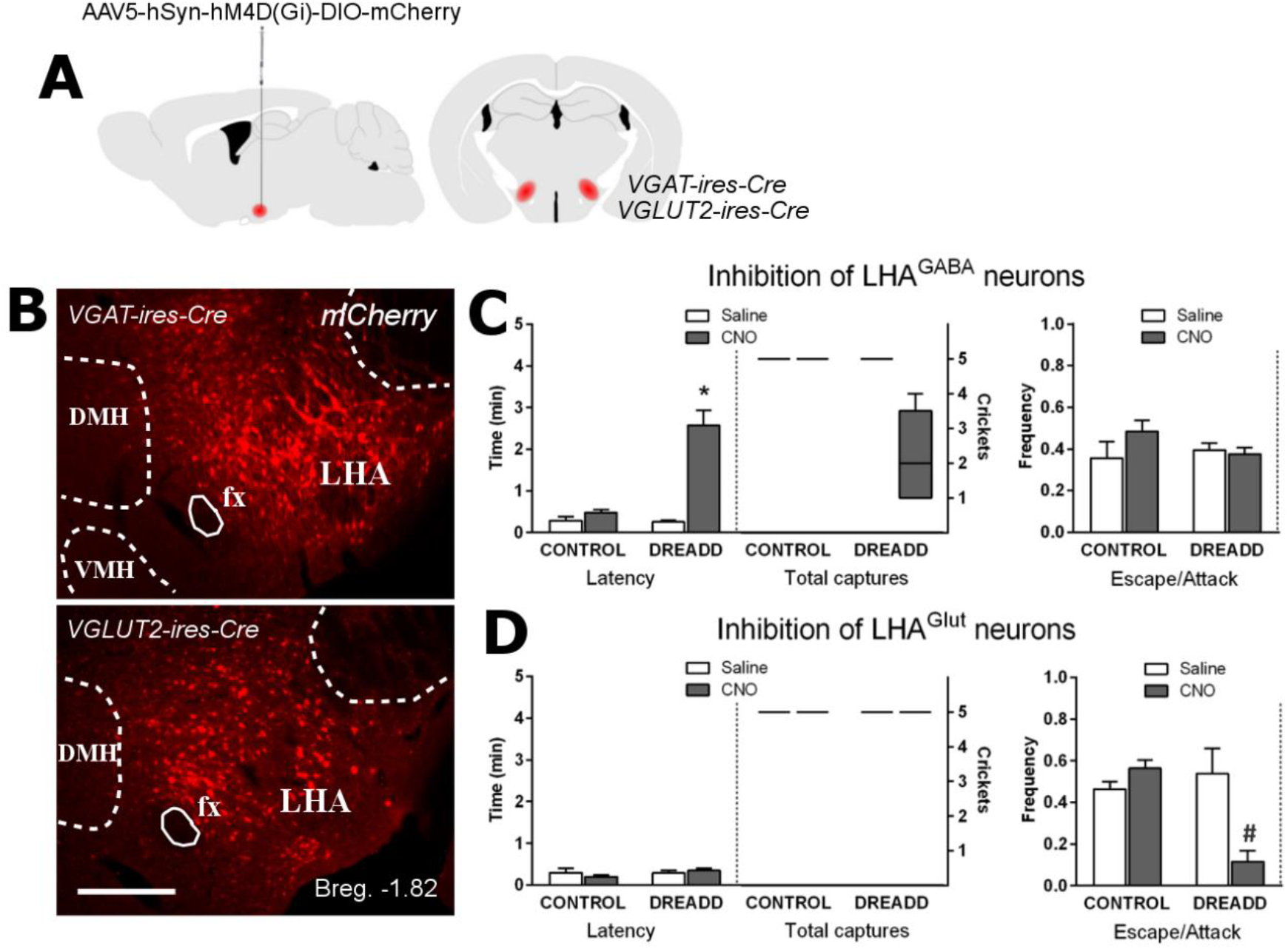
Functional role of LHA^GABA^ and LHA^Glut^ neurons on predatory hunting and social defensive behavior. **(A)** Schematics showing the bilateral injection in the LHA of Cre-dependent AAV expressing hM4D inhibitory DREADD in VGAT-IRES-Cre and VGLUT2-IRES-Cre mice. **(B)** Fluorescence photomicrographs showing neurons expressing mCherry (fused to HM4D) in the LHA of VGAT-IRES-Cre (top) and VGlut2-IRES-Cre (bottom) mice. DMH – dorsomedial hypothalamic nucleus; fx – fornix; LHA – lateral hypothalamic area; VMH – ventromedial hypothalamic nucleus. Scale bar, 150 μm. **(C)** The effect of CNO-induced inhibition of LHA^GABA^ neurons on predatory (latency to catch the first prey and total number of captures; left) and social defensive (escape/attack ratio; right) behaviors. Groups: CONTROL (*n* = 5); DREADD (*n* = 9). For the latency to catch the first prey, univariate ANOVA a significant effect for the factor virus (HM4D+ and control, F1,12= 9.04; p= 0.01) and the factor treatment (CNO and Saline, F1,12= 124.46; p<0.001), and a significant interaction between the factors (Virus x Treatment, F1,12= 39.9; p<0.001). Post hoc pairwise comparisons (Tukey’s HSD test) revealed that CNO treated HM4D+ animals presented a significant increase in the latency to catch the first prey (*p<0.001). CNO treated HM4D+ animals also reduced the number of captured crickets. For the escape/attack ratio during SD, univariate ANOVA revealed no significant effect for the factors virus (F1,12= 0.61; p=0.45) and treatment (F1,12=1.28; p=0.28). **(D)** The effect of CNO-induced inhibition of LHA^Glut^ neurons on predatory (latency to catch the first prey and total number of captures; left) and social defensive (escape/attack ratio; right) behaviors. Groups: CONTROL (*n* = 5); DREADD (*n* = 5). For the escape/attack ratio, univariate ANOVA revealed a significant effect for the factor virus (HM4D+ and control, F1,8= 6.75; p= 0.032) and the factor treatment (CNO and Saline, F1,8= 5.41; p= 0.048), and a significant interaction between the factors (Virus x Treatment, F1,8= 14.12; p= 0.005). Post hoc pairwise comparisons (Tukey’s HSD test) revealed that CNO treated HM4D+ animals presented a significant decrease in the escape/attack ratio (#p<0.01). For latency to start hunting, univariate ANOVA revealed no significant effect for the factors virus (F1,8= 0.98; p= 0.35) and treatment (F1,8= 0.04; p= 0.84). CNO treated HM4D+ animals also did not change the number of captured crickets. The data for latency and escape/attack ratio are shown as mean ± SEM, and the data for the number crickets captured are represented as box plot graphs. **See S5 – videos**.

## Discussion

Here, we addressed how the LPAG mediates the seemingly opposite behavioral choices of fleeing from a conspecific aggressor and prey hunting. We first found that LPAG neuronal populations responding to predatory hunting and social defensive responses appear to be partially differentiated. Our functional analysis in Fos DD-Cre mice revealed that CNO-induced inhibition of LPAG neurons activated in response to IP disrupted prey hunting but did not alter escape/attack ratio during SD. Conversely, CNO-induced inhibition of LPAG SD-responding neurons did not change predatory behavior parameters and led to a decrease in the escape/attack ratio during SD. Thus, this functional analysis supports the idea of diverse neural populations in the LPAG influencing the predatory and social-defensive responses. We further observed that the photoinhibition of LPAG -LHA pathway impaired IP but did not alter escape/attack ratio during SD. This suggests distinct paths from the LPAG to organized IP and SD.

To differentiate neuronal populations in LPAG responding to insect predation (IP) and social defeat (SD), we combined immunofluorescence for Fos protein (IF) and fluorescent *in situ* hybridization for c-*fos* mRNA (FISH) using sequential exposure to IP and SD. In response to the second stimulus (SD), we found 35% newly activated neurons. However, this number of newly activated neurons may have been underestimated because we sacrificed the animals 5 minutes prior to the peak of c-*fos* mRNA levels to minimize the detection of newly synthesized Fos protein from the second stimulus. The activation of partially distinct LPAG neuronal populations in response to IP and SD may reflect the contribution from the diverse sources of inputs to the LPAG that were differentially activated during IP or SD. Thus, during IP, we have a striking activation of the lateral part of the superior colliculus’ intermediate gray layer, which integrates critical sensory information to prey detection and projects to the LPAG (Mota-Ortiz et al., 2009; Furigo et al., 2010). Conversely, animals exposed to a conspecific aggressor present strong activation in the ventrolateral part of the ventromedial hypothalamic nucleus and the dorsolateral pre-mammillary nucleus (Motta et al., 2009) that represent relevant sources of inputs to the LPAG (Mota-Ortiz et al., 2009; Motta et al., 2009).

Our functional analysis using pharmacogenetic inhibition in Fos DD-Cre mice revealed a cleaner cut between the LPAG neural populations mediating IP and SD. Thus, CNO-induced inhibition of LPAG neurons activated in response to IP specifically impaired the IP behavioral parameters, but not the escape/attack ratio during SD. Conversely, the pharmacogenetic silencing of LPAG neurons activated in response to SD affected SD exclusively but not IP. Considering the large overlap between the neurons activated in IP and SD, we expect that the pharmacogenetic silencing would affect both responses regardless of the TMP time-dependent Cre stabilization for IP or SD. Thus, the results suggest that the non-overlap LPAG neural population differentially activated in response to IP or SD should mediate the behavioral responses measured in this study. The present observations raise an interesting question that the neuronal population of LPAG shared by IP and SD may conceivably mediate similar functional adjustments that may occur in both responses whereas the second population of neurons were activated explicitly during one type of behavior to mediate the specific behavioral outcomes.

The lateral hypothalamic area is one of the main targets of the LPAG (LHA; Mota-Ortiz et al., 2009), and we found that LPAG neurons activated during IP provide a denser projection to the LHA than those activated during SD. LPAG contains mostly glutamatergic neurons (Lein et al. 2007); in line with this view, we found that the vast majority of LPAG neurons activated during IP and SD co-expressed *VGLUT2*mRNA. Considering the predominantly excitatory nature of LPAG neurons activated during IP and SD, we employed an optogenetic projection-based silencing approach to examine the effect of the inhibition of the LPAG – LHA projection on predatory hunting and social defense. Accordingly, photoinhibition of the LPAG – LHA pathway impaired IP but did not change SD.

The LHA is known to control predatory hunting, and the LPAG glutamatergic ascending projection to the LHA would conceivably provide a motivational drive to the LHA on its influence on predatory hunting. Our findings agree with the idea that apart from organizing stereotyped responses, the PAG may exert more complex roles and provide a primal emotional tone to influence prosencephalic sites mediating complex behavioral responses (Motta et al., 2017). In line with this view, an opioid-dependent mechanism in the LPAG has been shown to switch the motivation from maternal care to prey hunting in lactating rats (Miranda-Paiva et al., 2003; Sukikara et al., 2006). In addition, LPAG cytotoxic lesions impair place preference induced by morphine (De Oliveira, 2015) thus suggesting an influence on the motivational drive to seek appetitive reward. Conversely, the ascending projection to the LHA from the LPAG glutamatergic neurons does not seem to be critical to modulate SD. In this case, descending paths from the LPAG are likely to be more involved in controlling escape responses during SD.

The LHA has been involved in controlling predatory hunting and evasion. LHA^GABA^ neurons promote feeding (Li et al. 2018; Jennings et al. 2013, 2015; O’Connor et al. 2015) and predatory hunting including searching, pursuing, attacking, and capturing behaviors toward a moving prey (Li et al. 2018). Importantly, studies using pharmacogenetic manipulations showed that LHA^GABA^ neurons mediate consummatory behaviors regardless of the caloric content or biological relevance of the consumed stimuli (Navarro et al. 2016). Conversely, LHA^Glut^ neurons may respond to aversive stimuli (Lazaridis et al., 2019) and control evasion by predicting imminent dangers (Li et al., 2018).

We found that LHA neuronal populations responding to predatory hunting and social defensive responses appear to be partially differentiated as seen by combining immunofluorescence for Fos protein and fluorescent *in situ* hybridization for c-*fos* mRNA in animals exposed sequentially to IP and SD. We next examined the GABAergic and glutamatergic nature of the LHA neurons activated during IP and SD. The results showed that the number of LHA^GABA^ activated neurons was significantly higher after exposure to IP compared to SD whereas the number of LHA^Glut^ activated neurons was comparable in both conditions. To address the behavioral role of LHA^GABA^ and LHA^Glut^ neurons on predatory hunting and social defensive responses, we performed a pharmacogenetic inhibition in VGAT-IRES-Cre and VGlut2-IRES-Cre mice to silence the activity of the LHA^GABA^ and LHA^Glut^ neurons selectively. Our functional analysis revealed that pharmacogenetic inhibition of LHA^GABA^ neurons impaired IP, increased the latency to start hunting, and decreased the total number of captured crickets; however, it did not alter evasion during SD. Conversely, pharmacogenetic inhibition of LHA^Glut^ neurons decreased the escape/attack ratio during SD but did not influence IP. Our results agree well with the literature and support the idea LHA^GABA^ neurons are involved in predatory hunting whereas the LHA^Glut^ neurons control evasion from aversive events. Li et al. (2018) showed that LHA^GABA^ neurons drive predation through projections to the ventral tegmental area and ventrolateral LPAG. Notably, a GABAergic pathway from the central amygdalar nucleus to the ventrolateral PAG also drives predatory hunting (Han et al., 2017). Moreover, ventrolateral PAG local neurons might be responsible for the evasion behavior evoked by activation of LHA^Glut^ neurons (Li et al, 2018), which also promotes aversive responses thorough a projection to the lateral habenula (Lazaridis et al., 2019).

Our study showed that the LPAG contains two segregated neural populations that separately control opposite behavioral choices of predatory hunting and evasion from a social attack. The LPAG and the LHA appear to provide a parallel glutamatergic output to control evasion in response to a social attack. LPAG control over predatory behavior involves an ascending glutamatergic pathway to the LHA that likely influences LHA^GABA^ neurons driving predatory hunting. Studies examining morphine-related influences on maternal care and place preference suggest that, apart from prey hunting, this glutamatergic pathway from the LPAG to the LHA may conceivably have a more widespread control on the motivational drive to seek appetitive rewards. Future studies are needed to address the general role of the LPAG as a player in rewarding motivational mechanisms.

## METHODS

### Animals

Adult male mice, C57BL/6J, Fos DD-Cre, VGAT-ires-Cre and VGLUT-ires-Cre – all bred onto a C57BL/6J genetic background – were individually housed under controlled temperature (23°-25°C) and 12h light cycle with free access to water and standard laboratory diet (Nuvilab-Quimtia, Brazil). At the time of the experiments, animals were 6-10 weeks old, weighting approximately 25-28 g. All experiments and conditions of animal housing were carried out under institutional guidelines [Colégio Brasileiro de Experimentação Animal (COBEA)] and were in accordance with the NIH Guide for the Care and Use of Laboratory Animals (NIH Publications No. 80-23, 1996). Procedures were previously approved by the Committee on the Care and Use of Laboratory Animals of the Institute of Biomedical Sciences of the University of São Paulo, Brazil (Protocol number 085/2012). Experiments were always planned to minimize the number of animals used and their suffering.

C57BL/6J mice (n = 61) were obtained from USP-local breeding facilities. Fos DD-Cre mice (n = 23), were developed (see Dillingham et al. 2019) and kindly provided by Dr Lisa Stowers Laboratory (Department of Neuroscience, The Scripps Research Institute, La Jolla, CA, USA). VGAT-ires-Cre (Slc32a1^tm2(cre)Lowl^/J, n = 14) and VGLUT2-ires-Cre (Slc17a6^tm2(cre)Lowl^/J, n = 10) knock-in mice were acquired from Jackson Laboratories (stocks #016962 and #016963).

### Viruses

For the functional analysis of neuronal populations in LPAG activated by IP and SD, Fos DD-Cre mice were injected bilaterally into the LPAG (Bregma AP −3.8, ML ±0.2, DV – 2.4) with a mixture of 20 nl of AAV5-hSyn-hM4D(Gi)-DIO-mCherry (titer ≥ 5×10^12^ vg/mL) and 20 nl of AAV5-hSyn-DIO-EYFP (titer ≥ 7×10^12^ vg/mL) (Dr. Bryan Roth, University of North Carolina’s Vector Core, NC, USA). For the functional analysis of LHA-projecting LPAG terminals, wild type mice were injected bilaterally into the LPAG with 40 nl of AAV5-hSyn-eArch3.0-EYFP (titer ≥ 4.4×10^12^ vg/mL) or AAV5-hSyn-EYFP-PA (titer ≥ 4.4×10^12^ vg/mL) (control virus) (Dr. Karl Deisseroth, University of North Carolina’s Vector Core, NC, USA) and optical fibers were implanted bilaterally into LHA (Bregma AP –1.55, ML ±1.1, DV –4.9). For the functional analysis of LHA^GABA^ and LHA^Glut^ neurons on predatory hunting and social defensive behavior, VGLUT2-ires-Cre and VGAT-ires-Cre animals were injected bilaterally into LHA (Bregma AP –1.55, ML ±1.1, DV –5.1) with a mixture of 50 nl of AAV5-hSyn-hM4D(Gi)-DIO-mCherry and 50 nl of AAV5-hSyn-DIO-EYFP (control virus).

### Stereotaxic surgery, viral injections and optical fiber implantation

Surgery for viral injection was performed at least 3 week before the first experimental day or the optical fiber implantation. During this time, animals remained undisturbed to recover from surgery and to allow viral expression. After the surgery for the implantation of optical fibers, animals remained undisturbed for one additional week to recover from surgery. For both viral injection and optic fiber implantation, mice were previously anesthetized in a box saturated with Isoforine (Cristália Laboratories, SP, Brazil) and then immediately head fixed on a stereotaxic instrument (Kopf Instruments, CA, USA). Anesthesia was maintained with 1-2% Isoforine/oxygen mix and body temperature was controlled with a heating pad. The scalp was incised and retracted, and the head position was adjusted to place bregma and lambda in the same horizontal plane. Viral vectors were injected with a 5 μl Hamilton Syringe (Neuros Model 7000.5 KH). Injections were delivered at a rate of 10 nl/min using a motorized pump (Harvard Apparatus). The needle was left in place for 5 minutes after each injection to minimize upward flow of viral solution after raising the needle. Optical fibers (Mono Fiber-optic Cannulae 200/230-0.48, Doric Lenses Inc. Quebeq, Canada) were implanted and fixed onto animal skulls with dental resin (DuraLay, IL, USA). After the behavioral tests, animals were perfused to verify injection sites by visualization of fluorescent markers.

### Behavioral procedures

#### Predatory hunting paradigm

The protocol used was adapted from Han et al (2018). Adult crickets (*Gryllus assimilis*) were used as preys. One week before the first experimental day animals were habituated to cricket hunting by presenting three intact crickets for two consecutive days. Food restriction was not required. All animals captured and consumed all crickets during each habituation. During the behavioral tests, five intact crickets were presented. Animals were let to hunt during 5 minutes after the first predatory attack. Behavior was recorded for subsequent analysis of the latency to catch the first prey the total number of crickets captured.

#### Resident-intruder paradigm

The protocol used was adapted from Miczek (1979). Adult Swiss male mice of approximately 14 weeks old, housed individually with sterilized but hormonally intact females, were used as residents. For the SD we used 8 residents, which were exposed only twice to intruders on the days of behavioral test. On the day of the behavioral test, the females were removed and replaced by intruders. The intruder was left interacting with the resident 5 minutes after the first attack. Behavior was recorded for subsequent analysis of the ratio of intruder escapes / resident attacks.

### Experimental design

#### Sequential exposure to predatory and social defensive behavior

The aim of this experiment was to identify IP- and SD-responding neuronal populations in LPAG by Fos/c-*fos* mRNA IF-FISH double labeling. First, C57BL/6J mice (n = 47) were used to determine the c-*fos* mRNA and Fos protein temporal dynamics in response to IP and SD. To this end, animals were sacrificed at basal time (n = 3), 30 min (n = 4), 1 hour (n = 4), 2 hours (n = 4) and 4 hours (n = 4) after the set of IP; and at basal time (n = 4), 10 min (n = 4), 20 min (n = 4), 30 min (n = 4), 1 hour (n = 4), 2 hours (n = 4) and 4 hours (n = 4) after the set of SD. Next, C57BL/6J mice (n = 18) were distributed into 4 experimental groups: *BASAL GROUP* (n = 4), animals received no treatment; *IP*+*IP GROUP* (n = 4), animals were exposed to IP (5 minutes), 1 hour and 40 minutes later re-exposed to IP (5 minutes), and after the second IP, were left for 10 minutes undisturbed before perfusion; *SD*+*SD GROUP* (n = 4), animals were exposed to SD (5 minutes), 1 hour and 40 minutes later re-exposed to SD (5 minutes), and after the second SD, were left for 10 minutes undisturbed before perfusion; *IP*+*SD GROUP* (n = 6), animals were exposed to IP (5 minutes), 1 hour and 40 minutes later exposed to SD (5 minutes), and after the SD, were left for 10 minutes undisturbed before perfusion. Groups of animals exposed repeatedly to the same treatment (IP+IP and SD+SD) were included to discard neuronal activation due to the mere manipulation of animals. Immediately after the treatments, animals were anesthetized and perfused. Brains were extracted and processed for the simultaneous detection of Fos protein and c-*fos* mRNA by IF-FISH double labeling.

#### Characterization of GABA-ergic and glutamatergic phenotypic nature of IP- and SD-responding neurons in LPAG and LHA

C57BL/6J mice (n = 13) were distributed into 3 experimental groups: *BASAL GROUP* (n=3), animals received no treatment; *IP GROUP* (n=5), animals were exposed to IP (5 minutes HC) and left for 10 minutes undisturbed before perfusion; and *SD GROUP* (n = 5), animals were exposed to SD (5 minutes) and left for 10 minutes undisturbed before perfusion. Animals were anesthetized and perfused after the treatments. Brains were extracted and processed for Fos*-VGAT* mRNA and Fos*-VGLUT2* mRNA IF-FISH double labeling in LPAG and LHA.

#### Pharmacogenetic inhibition of IP- and SD-responding populations

Fos DD-Cre mice (n = 23) were distributed into 2 experimental groups: *IP-Cre stabilized GROUP* (n=12), animals with Cre stabilization induced for predatory hunting; and *SD-Cre stabilized GROUP* (n=11), animals with Cre stabilization induced for social defensive behavior. Animals were previously injected bilaterally into LPAG with AAV5-hSyn-hM4D(Gi)-DIO-mCherry and AAV5-hSyn-DIO-EYFP vectors. After 3 weeks, animals were exposed to IP (*IP-Cre stabilized GROUP)* or SD (*SD-Cre stabilized GROUP*). 10 minutes after exposure, TMP was administrated intraperitoneally (i.p.,170 mg/kg) for stabilization of Cre expressed in response to IP or SD-active (Fos expressing) neurons. Animals were then kept undisturbed for 4 hours to allow stabilized Cre interacts with the viral vectors. After the procedure of Cre stabilization, animals remained undisturbed for 3 weeks to allow hM4D(Gi) expression before pharmacogenetic inhibition.

#### Optogenetic inhibition of LHA-projecting LPAG terminals

C57BL/6J mice (n = 10) were distributed into 2 experimental groups: *CONTROL GROUP* (n = 4), animals injected bilaterally into LPAG with an AAV vector not expressing eArch3.0 (AAV-hSyn-EYFP-PA); and *EARCH GROUP* (n = 6), animlas injected bilaterally into LPAG with an AAV vector expressing eArch3.0 (AAV5-hSyn-eArch3.0-EYFP). After 3 weeks, the animals received the optical fiber implantation, and remained undisturbed for 3 weeks before optogenetic inhibition.

#### Pharmacogenetic inhibition of LHA^GABA^ and LHA^Glut^ neurons

VGAT-ires-Cre mice (n = 14) were distributed into 2 experimental groups: *CONTROL GROUP* (n = 5), animals were injected bilaterally into LHA with a Cre-dependent AAV vector not expressing the inhibitory DREADD (AAV5-hSyn-DIO-EYFP); and *DREADD GROUP* (n = 9), animals were injected bilaterally into LHA with AAV5-hSyn-hM4D(Gi)-DIO-mCherry vector. VGLUT2-ires-Cre mice (n = 10) were distributed into 2 experimental groups: CONTROL GROUP (n = 5), animals were injected bilaterally into LHA with AAV5-hSyn-DIO-EYFP vector; and *DREADD GROUP* (n = 5), animals were injected bilaterally into LHA with AAV5-hSyn-hM4D(Gi)-DIO-mCherry vector. After surgery for viral injection, animals remained undisturbed for 3 weeks to allow viral expression before pharmacogenetic inhibition.

### Pharmacogenetic inhibition

Animals were previously habituated to handling and injections for the 3 days before the first experimental day (one habituation session per day). After the handling session, animals received 250 μl of saline i.p. On the first experimental day (day 1), animals (IP-Cre stabilized and SD-Cre stabilized groups for Fos DD-Cre animals; and CONTROL and DREADD groups for VGAT-ires-Cre and VGLUT2-ires-Cre animals) were injected with saline i.p. and 30 minutes later they were tested for predatory behavior. On the next day (day 2), clozapine-N-Oxide (CNO; Tocris Bioscience, UK) was injected i.p (1 mg/kg) for the pharmacogenetic inhibition of IP- and SD-responding neurons in Fos DD-Cre mice, LHA^GABA^ in VGAT-ires-Cre mice, and LHA^Glut^ in VGLUT2-ires-Cre mice, and 30 minutes later, animals were tested again for predatory behavior. After three days (day 6), animals of all groups were injected with saline and 30 minutes later they were tested for social defensive behavior. On the next day (day 7), animals were injected with CNO and 30 minutes later, animals were tested again for social defensive behavior. Behavioral responses were recorded during all experimental sessions for subsequent analysis.

### Optogenetic inhibition

For the optogenetic inhibition, animals were previously habituated to the optogenetic cables for the 3 consecutive days before the first experimental day (one habituation session per day). The habituation session consisted in plugging optogenetic cables to the implanted fiber-optic cannulae letting the animals explore their home cage for 5 minutes without any additional stimulus. The optogenetic inhibition was induced by exposing animals continuously to a yellow laser (589 nm, Low-Noise DPSS Laser System, Laserglow Techonologies). On the experimental day 1, animals (groups CONTROL and EARCH) were exposed to predatory behavior with the yellow laser turned OFF. On the experimental day 2, animals were exposed again to predatory behavior with the laser ON for the optogenetic inhibition of LHA terminals originating from LPAG. After three days (day 6), animals of both groups were exposed to social defensive behavior with the yellow laser turned OFF. On the next day (day 7), animals were animals were re-exposed to social defensive behavior with the laser ON. ON/OFF laser cycles were of 5 minutes for insect predation and 5 minutes for defensive behavior.

### Perfusion and histological processing

After the experimental procedures, animals were anesthetized in a box saturated with Isoforine and fixed by transcardial perfusion, introducing first a physiological saline solution (0.4% NaCl) and then a fixative solution (4% PFA + 3.8% Borax). After perfusion, brains were removed, post-fixed in the same solution and embedded in a cryoprotectant solution containing 30% sucrose in potassium phosphate-buffered saline (KPBS; 0.2 M NaCl, 43 mM potassium phosphate). Brains were then frozen in dry ice cooled isopentane at −50 °C, and preserved at −80 °C. For cryosectioning, brains were embedded in O.C.T. compound (Fisher Healthcare, USA) and sectioned using a Leica CM1850 cryostat. 20 μm-thick coronal sections were collected in cryoprotectant solution (0.05M sodium phosphate buffer pH 7.3, 30% ethyleneglycol, 20% glycerol) and stored at −20 °C.

### IF-FISH double labeling

#### Preparation and synthesis of RNA probes

c-*fos* DIG antisense riboprobe was *in vitro* transcribed (DIG RNA Labeling Mix, Roche) from an EcoRI fragment of c-*fos* cDNA (Dr Inder Verma, The Salk Institute, CA, USA; van Beveren et al. 1983), subcloned into pBluescript SK-1 (Stratagene). This construction was generously provided by Dr Antonio Armario (Universidad Autónoma de Barcelona, Barcelona, Spain). VGAT and VGLUT2 DIG antisense riboprobes were transcribed *in vitro* from purified PCR-amplified DNA fragments (PureLink Quick Gel Extraction Kit, Invitrogen, CA, USA). Forward and reverse primers used were 5’-GCCATTCAGGGCATGTTC-3’ and 5’-AGCAGCGTGAAGACCACC-3’ for VGAT DIG antisense, and 5’-CCAAATCTTACGGTGCTACCTC-3’ and 5’-TAGCCATCTTTCCTGTTCCACT-3’ for VGLUT2 DIG riboprobe. After *in vitro* transcription, the product was heated during 5 minutes at 65 °C. The probes were isolated through gel filtration columns (mini Quick Spin RNA Columns, Roche) and stored at −20°C.

#### IF-FISH procedure

The protocol for IF-FISH was adapted from Marin-Blasco et al. (2017). All the solutions used were previously treated with diethylpyrocarbonate (DEPC) and sterilized to ensure RNAse-free conditions. With the same purpose, glassware was previously baked at 200 °C and all consumables were sterile or pretreated with RNase Away (Sigma). For the IF, free-floating sections were washed with KPBS to remove the cryoprotectant solution and then incubated directly with polyclonal rabbit anti-c-Fos (PC-38; Calbiochem-Millipore) at 1:20000 in 0.5% Blocking Reagent (Roche) in KPBS for 16 h at 4 °C. Sections were then washed in KPBS and incubated with Anti-Rabbit Alexa 594 Goat IgG (H+L) (Invitrogen) at 1:500 during 2 h at random temperature (RT). Sections were then washed in KPBS and mounted on positive charged slides (Superfrost Plus, Thermo Scientific). FISH protocol was originally adapted from Simmons et al. (1989). Tissue was post-fixed in 4% PFA + 3.8% Borax, washed in KPBS and incubated for 15 minutes at 37 °C in presence of proteinase K (Roche) at 0.01 mg⁄mL in an appropriate buffer (0.1M Tris–HCl pH 8.0, 50mM EDTA pH 8.0). Sections were then rinsed in DEPC-treated water and acetylated during 10 minutes in 0.25% acetic anhydrous in 0.1 M TEA (triethanolamine) pH 8.0. Finally, sections were washed in 2X concentrated saline-sodium citrate solution (SSC), dehydrated through graded concentrations of ethanol and air-dried. Thereafter, 100 μL of hybridization buffer (50% formamide, 0.3 M NaCl, 10 mM Tris-Cl pH 8.0, 1 mM EDTA pH 8.0, 1X Denhardts, 10% dextrane sulphate, yeast tRNA 500 μg⁄mL and 10 mM DTT) containing the DIG-labeled probe (1:2000) were added onto each slide and covered with a glass coverslip. Sections were incubated for 16-18 h in a humid chamber at 60 °C. After this time, sections were washed in 4X SSC and RNA digested with RNase A (GE Healthcare) at 0.02 mg⁄mL in an appropriate buffer (0.5M NaCl, 10mM Tris–HCl pH 8.0, 1mM EDTA pH 8.0). After RNA digestion, sections were washed in descending concentrations of SSC (2X to 0.5X) containing 1mM DTT, and then heated at 60 °C in 0.1X SSC during 30 min. Sections were then equilibrated in Tris-buffered saline with Tween 20 (T-TBS; 0.1 M Tris-HCl pH 7.5, 0.15 M NaCl, 0.05% Tween 20) and incubated for 30 minutes in blocking buffer [1% bovine serum albumin (BSA) in TBS]. Blocking buffer was then removed and a peroxidase-conjugated anti-DIG antibody (anti-DIG-POD, Roche) was added at 1:2000 in 1% BSA in T-TBS. After incubation during 2 hours at RT, slides were washed in T-TBS. Signal was amplified using a tyramide signal amplification kit (TSA-plus Fluorescein, PerkinElmer). After the amplification, nuclei were counterstained with DAPI (Sigma) at 1:20000 in TBS and mounted using an aqueous based mounting medium (Fluoromount, Sigma). Slides were stored at 4°C in humid chambers until image capture.

### Image capture and analysis

Brain regions of interest were determined using a mouse brain stereotaxic atlas (Paxinos and Franklin, 2012). LPAG was captured and quantified at the level of the oculomotor nucleus (approximately between Bregma −3.79 mm and −4.23 mm). LHA was captured and quantified at the level of the ventromedial nucleus (approximately between Bregma −1.43 mm and −1.80 mm). Images were captured using an epi-fluorescence microscope (NIKON, Eclipse E400) coupled to a digital camera (NIKON, DMX 1200). ImageJ public domain image processing software (FIJI v1.47f) was used for image analysis.

#### IF–FISH Double Labeling Images

Background signal was determined by the mean integrated density (ID) value of 10 unlabeled regions of interest (ROIs) per projection. IF and FISH particles were considered positive cells when ID value of ROIs exceeded 3 standard deviations over the average background. This resulted in 3 different types of cells: positive only for protein (IF+/FISH−), positive only for mRNA (IF−/FISH+), and double labeled (IF+/FISH+). Average of at least 6 fields (one field per hemisphere of three slices) per brain area and animal was used for the statistical analysis. Images were coded before counting. The number of positive particles was corrected by the quantified area and density was expressed as the number of cells per mm^2^.

### Behavioral analysis

All behavioral sessions were recorded using a high-speed (120fps) camera (DMC-FZ200, Panasonic) and they were blindly scored using an ethological analysis software “The Observer” (version 5.0; Noldus Information Technology, Wageningen, The Netherlands). For predatory behavior, the following behavioral items were encoded: 1) the latency to capture the first prey; and 2) the total number of crickets captured. For social defensive behavior, behavioral items were: 1) escape or flight: number of flight episodes of the intruder such as running away and jumping; and 2) attacks: chasing and biting of the intruder by the resident. The ratio between the number of escapes and attacks was processed as the frequency of the escapes for each attack episode.

### Statistical analysis for cell counting

Statistical analysis was carried out with Statistical Package for Social Science software (SPSS, version 26 for Windows). Generalized linear model (GzLM; McCulloch and Searle 2001) was used to test between-subjects main effects. Significance is determined by the Wald chi-square statistic (χ^2^). After the main analysis, appropriate pair-wise comparisons were carried out and corrected by the Holm’s Sequential Bonferroni Procedure procedure (Holm, 1979). The criterion for significance was set at p<0.05. Particular comparisons were planned to avoid or reduce corrections for multiple comparisons inall tests. All samples to be statistically compared were processed in the same assay to avoid inter-assay variability.

### Statistical analysis for behavioral measurements

After testing for homogeneity of the variance with the Levene’s test, our behavioral data (latency to start hunting and escape / attack ratio) were logarithmic transformed. The data were analyzed using 2 × 2 univariate ANOVAs for each dependent variable (latency to start hunting and escape / attack ratio). The two factors (with two levels each) that entered the analysis were: time dependent Cre stabilization (during IP or SD) and treatment (CNO or saline) for the pharmacogenetic inhibition of neuronal populations in LPAG activated by IP and SD; virus (eArch+ and control) and treatment (laser on / off) for optogenetic inhibition of LHA-projecting LPAG terminals during IP and SD; and virus (hM4D+ and control) and treatment (CNO and saline) for the pharmacogenetic inhibition of GABAergic and glutamatergic LHA neurons during IP and SD. After obtaining a significant effect for each factor and a significant interaction between the factors, we applied Tukey’s HSD test post hoc analysis to isolate the respective effects. For the total number of captured crickets observed during IP experiments, we used box plot graphical tool to represent the variation of the observed data.

## Supporting information

S3_Fos DD Cre Predation CNO

S3_Fos DD Cre Predation Saline

S4_LPAG LHA Photoinhibition Laser OFF

S4_LPAG LHA Photoinhibition Laser ON

S5_vGAT LHA Predation CNO

S5_vGAT LHA Predation Saline

## Acknowledgments

This research was supported by Fundação de Amparo à Pesquisa do Estado de São Paulo (FAPESP) Research Grants #2014/05432-9, 2016/18667-0, 2017/04830, and 2017/09753-2.

## Notes

### Competing Interest Statement

The authors have declared no competing interest.

## References

Armbruster, B. N., Li, X., Pausch, M. H., Herlitze, S. & Roth, B. L. Evolving the lock to fit the key to create a family of G protein-coupled receptors potently activated by an inert ligand. Proc. Natl. Acad. Sci. U. S. A. 104, 5163–5168 (2007).

Assareh, N., Sarrami, M., Carrive, P. & McNally, G. P. The organization of defensive behavior elicited by optogenetic excitation of rat lateral or ventrolateral periaqueductal gray. Behav. Neurosci. 130, 406–414 (2016).

Bandler, R. & Shipley, M. T. Columnar organization in the midbrain periaqueductal gray: modules for emotional expression? Trends Neurosci. 17, 379–89 (1994).

Besson, J.-M., Fardin, V. & Olivéras, J.-L. Analgesia Produced by Stimulation of the Periaqueductal Gray Matter: True Antinoceptive Effects Versus Stress Effects. in The Midbrain Periaqueductal Gray Matter 121–138. Springer US (1991).

Cezario, A. F., Ribeiro-Barbosa, E. R., Baldo, M. V. C. & Canteras, N. S. Hypothalamic sites responding to predator threats - The role of the dorsal premammillary nucleus in unconditioned and conditioned antipredatory defensive behavior. Eur. J. Neurosci. 28, 1003–1015 (2008).

Comoli, E., Ribeiro-Barbosa, E. R. & Canteras, N. S. Predatory hunting and exposure to a live predator induce opposite patterns of Fos immunoreactivity in the PAG. Behav. Brain Res. 138, 17–28 (2003).

de Andrade Rufino, R., Mota-Ortiz, S. R., De Lima, M. A. X., Baldo, M. V. C. & Canteras, N. S. The rostrodorsal periaqueductal gray influences both innate fear responses and acquisition of fear memory in animals exposed to a live predator. Brain Struct. Funct. 224, 1537–1551 (2019).

Deng, H., Xiao, X. & Wang, Z. Periaqueductal gray neuronal activities underlie different aspects of defensive behaviors. J. Neurosci. 36, 7580–7588 (2016).

De Oliveira, W.F. Analysis of the rotrolateral portion of the periaqueductal gray (PAGrl) in drug seeking behavior. Ph.D. Thesis. Institute of Biomedical Sciences. University of São Paulo. pp. 86 (2015).

Depaulis, A., Keay, K. A. & Bandler, R. Longitudinal neuronal organization of defensive reactions in the midbrain periaqueductal gray region of the rat. Exp. Brain Res. 90, 307–318 (1992).

Di Scala, G., Mana, M. J., Jacobs, W. J. & Phillips, A. G. Evidence of Pavlovian conditioned fear following electrical stimulation of the periaqueductal grey in the rat. Physiol. Behav. 40, 55–63 (1987).

Di Scala, G. & Sandner, G. Conditioned place aversion produced by microinjections of semicarbazide into the periaqueductal gray of the rat. Brain Res. 483, 91–97 (1989).

Dillingham, B. C. et al. Fear Learning Induces Long-Lasting Changes in Gene Expression and Pathway Specific Presynaptic Growth. bioRxiv 571331 (2019)

Fanselow, M. S. The Midbrain Periaqueductal Gray as a Coordinator of Action in Response to Fear and Anxiety. in The Midbrain Periaqueductal Gray Matter 151–173. Springer US (1991).

Furigo, I. C. et al. The role of the superior colliculus in predatory hunting. Neuroscience 165, 1–15 (2010).

Gruber-Dujardin, E. Role of the periaqueductal gray in expressing vocalization. in Handbook of Behavioral Neuroscience vol. 19, 313–327 (2010).

Han, W. et al. Integrated Control of Predatory Hunting by the Central Nucleus of the Amygdala. Cell 168, 311–324.e18 (2017).

Holm, S. A simple sequentially rejective multiple test procedure. Scand. J. Stat. 6, 65–70 (1979).

Iwamoto, M., Björklund, T., Lundberg, C., Kirik, D. & Wandless, T. J. A general chemical method to regulate protein stability in the mammalian central nervous system. Chem. Biol. 17, 981–988 (2010).

Jennings, J. H. et al. Visualizing hypothalamic network dynamics for appetitive and consummatory behaviors. Cell 160, 516–527 (2015).

Jennings, J. H., Rizzi, G., Stamatakis, A. M., Ung, R. L. & Stuber, G. D. The Inhibitory Circuit Architecture of the Lateral Hypothalamus Orchestrates Feeding. Science (80) 341, 1517–1521 (2013).

Jürgens, U. The role of the periaqueductal grey in vocal behaviour. Behav. Brain Res. 62 107–117 (1994).

Kim, E. J. et al. Dorsal periaqueductal gray-amygdala pathway conveys both innate and learned fear responses in rats. Proc. Natl. Acad. Sci. U. S. A. 110, 14795–14800 (2013).

Kincheski, G. C., Mota-Ortiz, S. R., Pavesi, E., Canteras, N. S. & Carobrez, A. P. The Dorsolateral Periaqueductal Gray and Its Role in Mediating Fear Learning to Life Threatening Events. PLoS One 7, e50361 (2012).

Kovács, K. J. & Sawchenko, P. E. Sequence of stress-induced alterations in indices of synaptic and transcriptional activation in parvocellular neurosecretory neurons. J. Neurosci. 16, 262–273 (1996).

Kovács, K. J. Measurement of immediate-early gene activation-c-fos and beyond. J. Neuroendocrinol. 20, 665–672 (2008).

Lazaridis, I. et al. A hypothalamus-habenula circuit controls aversion. Mol. Psychiatry 24, 1351–1368 (2019).

Lein, E. S. et al. Genome-wide atlas of gene expression in the adult mouse brain. Nature 445, 168–176 (2007).

Li, Y. et al. Hypothalamic Circuits for Predation and Evasion. Neuron 97, 911–924.e5 (2018).

Lonstein, J. S. & Stern, J. M. Role of the midbrain periaqueductal gray in maternal nurturance and aggression: C-fos and electrolytic lesion studies in lactating rats. J. Neurosci. 17, 3364–3378 (1997).

Lonstein, J. S. & Stern, J. M. Site and behavioral specificity of periaqueductal gray lesions on postpartum sexual, maternal, and aggressive behaviors in rats. Brain Res. 804, 21–35 (1998).

Lovick, T. A. Integrated activity of cardiovascular and pain regulatory systems: Role in adaptive behavioural responses. Progress in Neurobiology 40, 631–644 (1993).

McCulloch C. E. Searle S.R. Generalized, linear, and mixed models. New York: John Wiley & Sons (2001).

Marín-Blasco, I., Muñoz-Abellán, C., Andero, R., Nadal, R. & Armario, A. Neuronal Activation After Prolonged Immobilization: Do the Same or Different Neurons Respond to a Novel Stressor? Cereb. Cortex 28, 1233–1244 (2018).

Miczek, K. A. A new test for aggression in rats without aversive stimulation: Differential effects of d-amphetamine and cocaine. Psychopharmacology (Berl). 60, 253–259 (1979).

Miranda-Paiva, C. M., Ribeiro-Barbosa, E. R., Canteras, N. S. & Felicio, L. F. A role for the periaqueductal grey in opioidergic inhibition of maternal behaviour. Eur. J. Neurosci. 18, 667–674 (2003).

Mota-Ortiz, S. R. et al. The periaqueductal gray as a critical site to mediate reward seeking during predatory hunting. Behav. Brain Res. 226, 32–40 (2012).

Mota-Ortiz, S. R., Sukikara, M. H., Felicio, L. F. & Canteras, N. S. Afferent connections to the rostrolateral part of the periaqueductal gray: A critical region influencing the motivation drive to hunt and forage. Neural Plast. (2009).

Motta, S. C. et al. Dissecting the brain’s fear system reveals the hypothalamus is critical for responding in subordinate conspecific intruders. Proc. Natl. Acad. Sci. U. S. A. 106, 4870–4875 (2009).

Motta, S. C., Carobrez, A. P. & Canteras, N. S. The periaqueductal gray and primal emotional processing critical to influence complex defensive responses, fear learning and reward seeking. Neurosci. Biobehav. Rev. 76, 39–47 (2017).

Navarro, M. et al. Lateral hypothalamus GABAergic neurons modulate consummatory behaviors regardless of the caloric content or biological relevance of the consumed stimuli. Neuropsychopharmacology 41, 1505–1512 (2016).

O’Connor, E. C. et al. Accumbal D1R Neurons Projecting to Lateral Hypothalamus Authorize Feeding. Neuron 88, 553–564 (2015).

Paxinos G., Franklin K.B.J. The Mouse Brain in Stereotaxic Coordinates, 4rd edn, Academic Press (2012).

Pobbe, R. L. H., Zangrossi, H., Blanchard, D. C. & Blanchard, R. J. Involvement of dorsal raphe nucleus and dorsal periaqueductal gray 5-HT receptors in the modulation of mouse defensive behaviors. Eur. Neuropsychopharmacol. 21, 306–315 (2011).

Sakuma, Y. & Pfaff, D. W. Facilitation of female reproductive behavior from mesencephalic central gray in the rat. Am. J. Physiol. - Regul. Integr. Comp. Physiol. 6, (1979).

Sando, R. et al. Inducible control of gene expression with destabilized Cre. Nat. Methods 10, 1085–1088 (2013).

Simmons, D. M., Arriza, J. L. & Swanson, L. W. A complete protocol for in situ hybridization of messenger RNAs in brain and other tissues with radiolabeled single-stranded RNA probes. J. Histotechnol. 12, 169–181 (1989).

Souza, R. R. & Carobrez, A. P. Acquisition and expression of fear memories are distinctly modulated along the dorsolateral periaqueductal gray axis of rats exposed to predator odor. Behav. Brain Res. 315, 160–167 (2016).

Sukikara, M. H., Mota-Ortiz, S. R., Baldo, M. V., Felício, L. F. & Canteras, N. S. A role for the periaqueductal gray in switching adaptive behavioral responses. J. Neurosci. 26, 2583–2589 (2006).

Sukikara, M. H., Mota-Ortiz, S. R., Baldo, M. V., Felicio, L. F. & Canteras, N. S. The periaqueductal gray and its potential role in maternal behavior inhibition in response to predatory threats. Behav. Brain Res. 209, 226–233 (2010).

Tryon, V. L. & Mizumori, S. J. Y. A novel role for the periaqueductal gray in consummatory behavior. Front. Behav. Neurosci. 12, (2018).

Wang, L. et al. Hypothalamic Control of Conspecific Self-Defense. Cell Rep. 26, 1747–1758.e5 (2019).

Zangenehpour, S. & Chaudhuri, A. Differential induction and decay curves of c-fos and zif268 revealed through dual activity maps. Mol. Brain Res. 109, 221–225 (2002).

