## Supplementary Data and Material for "The lateral periaqeductal gray and its role in controlling the opposite behavioral choices of predatory hunting and social defense"

**S1. c-*fos* mRNA and Fos protein temporal dynamics in response to IP and SD.**

To determine the optimum sacrificing time points, we quantified the number of Fos protein and c-*fos* mRNA positive neurons in LPAG at different time points after IP and SD. To differentiate neuronal populations in LPAG responding to insect predation (IP) and social defeat (SD), we exposed animals first to IP and subsequently to SD, particularly considering that exposure to a stressing stimulus such as SD would affect animals' motivation for hunting. The aim of this temporal profile was to find the best sacrificing time for animals exposed sequentially to IP and SD. Therefore, we performed the c-*fos* mRNA and Fos protein temporal dynamics to determine the time points when would have the maximum Fos protein expression with imperceptible levels of c-*fos* mRNA in response to the IP (the first stimulus), and optimum c-*fos* mRNA levels with reduced levels of Fos protein in response to the SD (the second stimulus). 2 hours after the set of IP, we found maximum levels of Fos protein levels and imperceptible c-*fos* mRNA signal; and 20 minutes after to the set of SD, we obtained maximum c-*fos* mRNA levels (Figure S1). However, low but perceptible synthesized Fos protein emerged at this time point in response to SD (Figure S1). Thus, we decided to shorten this time point, and sacrificed the animals 15 minutes after the set of SD when c-*fos* mRNA levels are close to the maximum, and less significant Fos protein levels were detected.

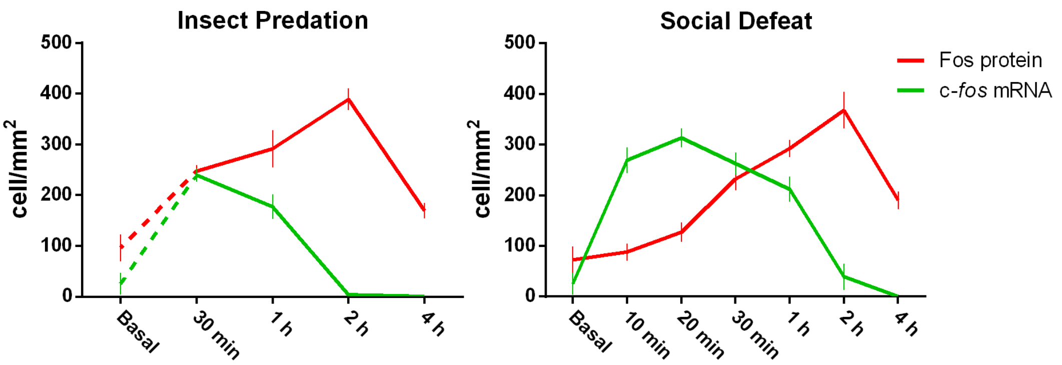


**Supplementary Figure 1. c-*fos* mRNA and Fos protein temporal dynamics in response to IP and SD.** Curves representing the c-*fos* mRNA and Fos protein temporal dynamics in response to IP and SD. C57BL/6J mice (n = 47) were used to determine the c-*fos* mRNA and Fos protein temporal dynamics in response to IP and SD. To this end, animals were sacrificed at basal time (n = 3), 30 min (n = 4), 1 hour (n = 4), 2 hours (n = 4) and 4 hours (n = 4) after the set of IP; and at basal time (n = 4), 10 min (n = 4), 20 min (n = 4), 30 min (n = 4), 1 hour (n = 4), 2 hours (n = 4) and 4 hours (n = 4) after the set of SD. Values expressed as means ± SEM of cells per $mm^2$.

### S2. Phenotypic characterization of the LPAG neurons activated in the SD and IP.

To characterize the GABAergic or glutamatergic nature of LPAG neurons activated during IP and SD, we applied an IF-FISH double labeling for Fos protein and mRNAs for VGAT and VGLUT2. The results showed that the vast majority of LPAG neurons activated during IP and SD co-expressed *VGLUT2*mRNA, and just an exceedingly small number was *VGAT* mRNA positive (Figure S2). Notably, Group SD presented a lower number of Fos/*VGLUT2* mRNA neurons when compared to IP group ($p < 0.01$) (Figure S2E). The present results are in line with previous observations showing that the LPAG contains mostly glutamatergic neurons (Lein et al. 2007) and support the idea that LPAG neurons activated during SD and IP should provide glutamatergic projections to their projection targets.

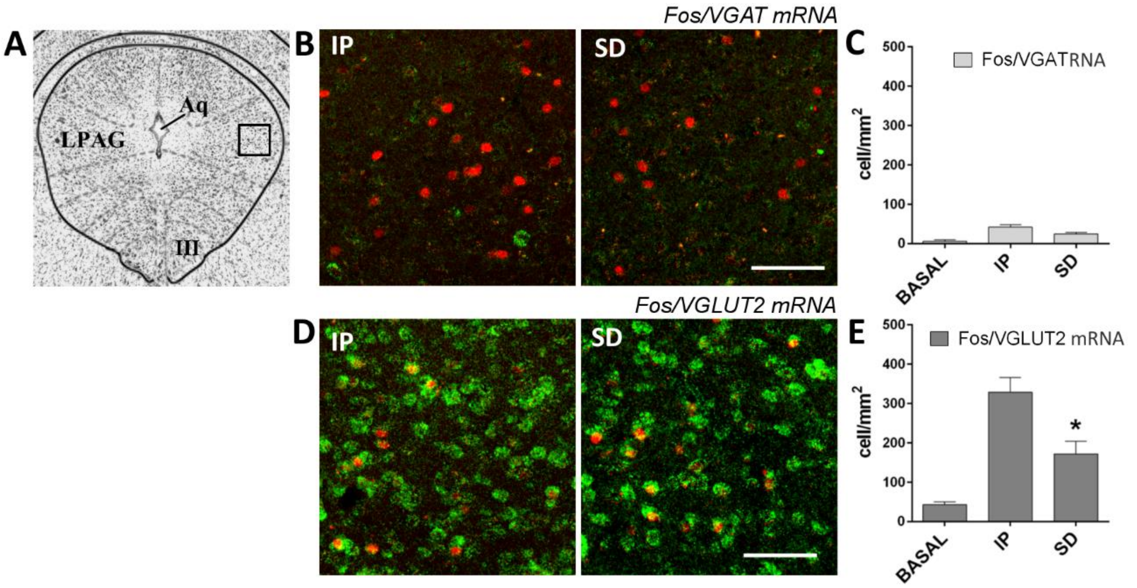


**Supplementary Figure 2. Analysis of the GABAergic and glutamatergic nature of LPAG neurons recruited in response to predatory hunting and social defensive behavior.**

(A) Photomicrograph of a representative transverse thionin-stained section of the LPAG at the level of the oculomotor nucleus (III). Square-delineated region indicates the approximate location of higher magnification fluorescence photomicrographs showing FOS / VGAT and VGLUT2 mRNAs double labeling in (B) and (D). Aq – cerebral aqueduct; III – oculomotor nucleus

(B) Representative fluorescence photomicrographs illustrating *Fos/VGAT* mRNA double labeling in LPAG of animals exposed to IP and SD. Fos protein positive cells are labeled by immunofluorescence in red (Alexa 594) and *VGAT* mRNA positive cells are labeled by fluorescent *in situ* hybridization in green (FITC). Scale bar, 75 μm.

(C) Median values of the density of Fos/VGATmRNA double labeled cells in the BASAL ($n = 3$), IP ($n = 6$), and SD ($n = 6$) groups.

(D) Representative fluorescence photomicrographs illustrating *Fos/VGULT2* mRNA double labeling in LPAG of animals exposed to IP and SD. Fos protein positive cells are labeled by immunofluorescence in red (Alexa 594) and *VGLUT2* mRNA positive cells are labeled by fluorescent *in situ* hybridization in green (FITC). Scale bar, 75 μm.

(E) Median values of the density of Fos/VGLUT2mRNA double labeled cells in the BASAL ($n = 3$), IP ($n = 6$), and SD ($n = 6$) groups. GzLM analysis of the number of Fos/*VGLUT2* mRNA neurons showed a significant effect of group factor (groups Basal, IP, and SD) [$\chi^2(2) = 44.83$, $p < 0.001$]. ***** indicates significant differences in *Fos-VGLUT2* mRNA neurons versus IP group ($p < 0.01$).

**S3. Videos – Functional analysis of neuronal populations in LPAG activated by IP and SD.** Videos of Fos DD-Cre mice expressing inhibitory DREADD in LPAG neurons activated during IP or SD tested during prey hunting (saline-treated - Fos DD-Cre Predation Saline; and CNO-treated - Fos DD-Cre Predation CNO).

**S4. Videos – Functional analysis of LHA-projecting LPAG terminals during IP and SD.** Videos of C57BL/6 mice expressing eArch3.0 in LPAG projecting fibers to the LHA that receive photoinhibition during prey hunting (Laser Off – LPAG-LHA Photoinhibition Laser OFF and Laser On – LPAG-LHA Photoinhibition Laser ON).

**S5. Videos – Functional role of $LHA^{GABA}$ neurons on predatory hunting.**

Videos comparing VGAT-IRES-Cre mice expressing inhibitory DREADD or EYFP (control group) in $LHA^{GABA}$ neurons tested during prey hunting (saline-treated – VGAT LHA Predation Saline; and CNO-treated – VGAT LHA Predation CNO).
